# *ovo^D^* co-selection: a method for enriching CRISPR/Cas9-edited alleles in *Drosophila*

**DOI:** 10.1101/310854

**Authors:** Ben Ewen-Campen, Norbert Perrimon

## Abstract

Screening for successful CRISPR/Cas9 editing events remains a time consuming technical bottleneck in the field of *Drosophila* genome editing. This step can be particularly laborious for events that do not cause a visible phenotype, or those which occur at relatively low frequency. A promising strategy to enrich for desired CRISPR events is to co-select for an independent CRISPR event that produces an easily detectable phenotype. Here, we describe a simple negative co-selection strategy involving CRISPR-editing of a dominant female sterile allele, *ovo^D1^*. In this system (“*ovo^D^* co-selection”), the only functional germ cells in injected females are those that have been edited at the *ovo^D1^* locus, and thus 100% of the offspring of these flies have undergone editing of at least one locus. We demonstrate that *ovo^D^* co-selection can be used to enrich for knock-out mutagenesis via nonhomologous end-joining (NHEJ), and for knock-in alleles via homology-directed repair (HDR). Altogether, our results demonstrate that *ovo^D^* co-selection reduces the amount of screening necessary to isolate desired CRISPR events in *Drosophila*.

## INTRODUCTION

In the five short years since CRISPR/Cas9-based genome-editing was first demonstrated in *Drosophila* (Bassett *et al*. 2013; Ren *et al*. 2013; Gratz *et al*. 2014; Port *et al*. 2014), the technique has revolutionized fruit fly research, just as it has for nearly every organism studied (reviewed in Sternberg and Doudna 2015). Because CRISPR/Cas9 generates targeted double-stranded breaks in DNA, this technique can be used to create both “knock-out” mutations, via imprecise repair of Cas9-induced lesions via the non-homologous end-joining pathway (NHEJ), as well as “knock-in” mutations, where an exogenously-supplied DNA donor serves as a template for homology-directed repair (HDR) (Gratz *et al*. 2014). Indeed, a number of genome-wide *Drosophila* collections are currently being generated for both knock-outs (Kondo *et al*. 2017), and knock-ins (e.g. Lee *et al*. 2018).

Despite the enormous power of CRISPR/Cas9 for genome editing, screening for successful genome-editing events remains a time-consuming and laborious technical bottleneck in all organisms and in cell culture. In response to this challenge, a number of techniques have been developed to enrich and/or select for desired CRISPR events, collectively referred to as “CRISPR co-selection” (aka “co-CRISPR” or “CRISPR coconversion”) (Kim *et al*. 2014; Arribere *et al*. 2014; Liao *et al*. 2015; Shy *et al*. 2016; Ge *et al*. 2016; Agudelo *et al*. 2017). CRISPR co-selection is based on the observation that when two independent short guide RNAs (sgRNAs) and Cas9 protein are introduced to a population of cells simultaneously, CRISPR events tend to cooccur at both loci within individual cells at a higher-than-random frequency. CRISPR co-selection exploits this observation by introducing an sgRNA targeting a marker locus that produces an easily detectable and/or selectable phenotype, together with an sgRNA targeting the gene-of-interest. Successful variations on this strategy have been developed for *C. elegans* (Kim *et al*. 2014; Arribere *et al*. 2014), *Drosophila* (Ge *et al*. 2016; Kane *et al*. 2017), and for mammalian cell culture (Liao *et al*. 2015; Shy *et al*. 2016; Agudelo *et al*. 2017).

In *Drosophila*, the most common technique for generating CRISPR/Cas9 germ line mutations involves injecting a plasmid that encodes a U6-driven sgRNA (along with an HDR donor constructs, in the case of a knock-in) into embryos that express Cas9 in their germline (Port *et al*. 2015). As injected embryos develop, CRISPR/Cas9 editing occurs in a subset of each embryo’s germ cells, resulting in adult flies with mosaic germ line stem cells. Once mature, these injected flies are out-crossed, and their offspring are screened for successful editing events. While this strategy is broadly effective, the screening step remains particularly laborious for target loci whose disruption does not cause a visible phenotype, and/or for sgRNAs with low editing efficiency. Thus, methods to enrich for desired CRISPR/Cas9 events would greatly aid the rapidly growing field of *Drosophila* genome editing.

Here, we describe a simple CRISPR enrichment strategy where the co-selected phenotype is female fertility itself. This system is based on rescuing a fully penetrant dominant female sterile allele, *ovo^D1^* (Busson *et al*. 1983), using CRISPR/Cas9 genome editing. In this strategy, co-editing of the *ovo ^D1^* allele rescues germ cells that would otherwise be fully non-functional, and therefore 100% of eggs laid have necessarily undergone editing of at least one locus. Thus, unlike the two previously described co-selection strategies based on coselection for the visible markers *ebony* or *white* (Ge *et al*. 2016; Kane *et al*. 2017), our method simply removes from the population any germ cell that has not undergone editing of at least one locus. We show that this method, which we term *“ovo°* co-selection” successfully enriches for both knock-outs and knock-ins, and thus simplifies the screening step required for the generation of CRISPR mutations in *Drosophila*.

## MATERIALS AND METHODS

### sgRNA cloning and preparation

All sgRNA sequences are given in Table S1. sgRNAs targeting *ovo^D1^* were designed using the *Drosophila Resource Screening Center Find CRISPR v2* online tool (http://www.flyrnai.org/crispr2/), then independently screened for potential off-targets using the CRISPR Optimal Target Finder tool (http://tools.flycrispr.molbio.wisc.edu/targetFinder/index.php). Sources for additional sgRNAs are given in Table S1. sgRNAs were cloned into the pCFD3 vector as described (Port *et al*. 2014). sgRNA plasmids were purified using QIAprep miniprep kit (QIAGEN), then prepared for injection as follows: either single sgRNAs or pooled sgRNAs were purified using a fresh mini-prep column (QIAGEN), washed twice with Buffer PB, once with Buffer PE, then eluted in injection buffer. For initial characterization of the *ovo^D^* coconversion using *ebony*, 4 μg of sgRNA-*ovo^D1^* and 4μg of sgRNA-ebony plasmid were pooled, purified as described above, and eluted in 50 μL of standard *Drosophila* injection buffer. For subsequent *ebony* coselection experiments, 1.25 μg of sgRNA-*ovo^D1^* and 2.5 μg of sgRNA-ebony were pooled and purified in 20 μL of injection buffer. For knock-in experiments, 1 μg of sgRNA-*ovo^D1^*, 2 μg of sgRNA-target-gene, and 3 μg of HDR donor plasmid were pooled and purified as above, then eluted in 20μL of injection buffer.

### Fly work

*Drosophila* were maintained on a standard cornmeal diet, and crosses were maintained at either 25°C or 27°C, always consistent within a given experiment. *ovo^D1^* (K1237) (Busson *et al*. 1983) flies are kept as attached-X stocks, composed of C(1)DX,*y f/Y* females and *ovo^D1^ /Y* males. Table S2 lists all genotypes used in this study. To generate *ovo^D1^;; nos-Cas9* embryos for injection, male *ovo^D1^* flies were crossed to female *yv;; nos-Cas9^attP2^* (Ren *et al*. 2013) in bottles, then transferred to grape juice plates for embryo collections. Injections were performed following standard procedures, using sgRNA concentrations given below. Any injection where ≤ 5 G0s of either sex was obtained was discarded.

### Scoring fertility, mutant alleles, and knock-in efficiency

Injected G0 flies were mated individually to two opposite-sex flies (of various genotype depending on the gene to be scored) in vials of standard food supplemented with yeast powder, then flipped to fresh vials after four to five days later. Any fly that did not produce any offspring was scored sterile. To screen for *ebony* alleles, injected G0 flies were crossed to balancer lines containing independent *ebony* mutations (either *w;; Ly / TM6b Tb* or *w;; TM3 Sb / TM6b Tb*), and the proportion of phenotypically *ebony* flies was scored for each individual G0 cross. To screen for knock-ins, RFP+ or GFP+ eyes were scored at the adult stage using a fluorescent dissecting scope.

### Allele sequencing

To analyze the sequence of mutant alleles, genomic DNA was extracted from single flies by homogenizing flies in 50-100 μL of DNA extraction buffer (10mM Tris-Cl pH 8.2, 1mM EDTA, 25mM NaCL, 200 μg/mL Proteinase K), incubating at 37°C for 20-30 minutes, then boiling at 98°C for ~90 seconds. 1 μL of genomic DNA was used as template in a 20 μL PCR reaction amplifying a fragment that includes the targeted region (670 bp for *ebony*, F primer = ATCCTTGGTCACTGCCTTGG, R primer = CTATCAGCCCAGCACTACGG) using Phusion High Fidelity polymerase (New England BioLabs). PCR products were purified using a QIAquick PCR purification kit (QIAGEN) or Exo-SAP-IT (Thermo), then Sanger sequenced at the Dana Farber/Harvard Cancer Center DNA sequencing facility (sequencing primer = CCATAGCTCCGCAATCGAGT.) The sequencing trace files, which represent a mixture of a wildtype allele and a mutant allele, were deconvoluted using Poly Peak Parser (http://yosttools.genetics.utah.edu/PolyPeakParser/).

### Statistical and graphical analysis

Paired t-tests were used to compare the proportion of founders amongst female G0s versus male G0s across all experiments in this study, and the proportion of mutant offspring per fertile G0 female versus fertile G0 male across all experiments in this study. Statistical analysis and graphing was conducted using Prism 7 (GraphPad Software.)

### Data Availability Statement

All fly strains and plasmids used in this are available from the authors upon request, and/or from the Drosophila Bloomington Stock Center and Addgene, respectively. The sgRNAs used in this study are described in Table S1. The fly stocks used in this study are described in Table S2. The authors affirm that all data necessary for confirming the conclusions within this article are present within the article, figures, and tables.

## RESULTS & DISCUSSION

### CRISPR/Cas9 editing of *ovo^D1^* restores function in female germ cells

The *ovo* gene encodes an X-linked transcription factor required for germline development and function specifically in female *Drosophila* (Busson *et al*. 1983; Perrimon 1984; Oliver *et al*. 1987). The *ovo^D1^* mutation is a single A>T base pair substitution in the second exon of *ovo* that introduces a novel start codon, generating a dominant negative form of the protein which causes 100% sterility in heterozygous *ovo^D1^/* + females (Mével-Ninio *et al*. 1996) (**Figure 1A**), with an observed 0.05% rate of spontaneous reversion in females (Busson *et al*. 1983; Perrimon and Gans 1983). However, if the *ovo^D1^* mutant allele is removed from germ cells during early development, for example via mitotic recombination, germ cell function can be restored (Perrimon 1984). This unique property of the *ovo^D1^* allele has led to its widespread use for generating homozygous germline clones (Chou and Perrimon 1996; Griffin *et al*. 2014).

**Figure 1.**
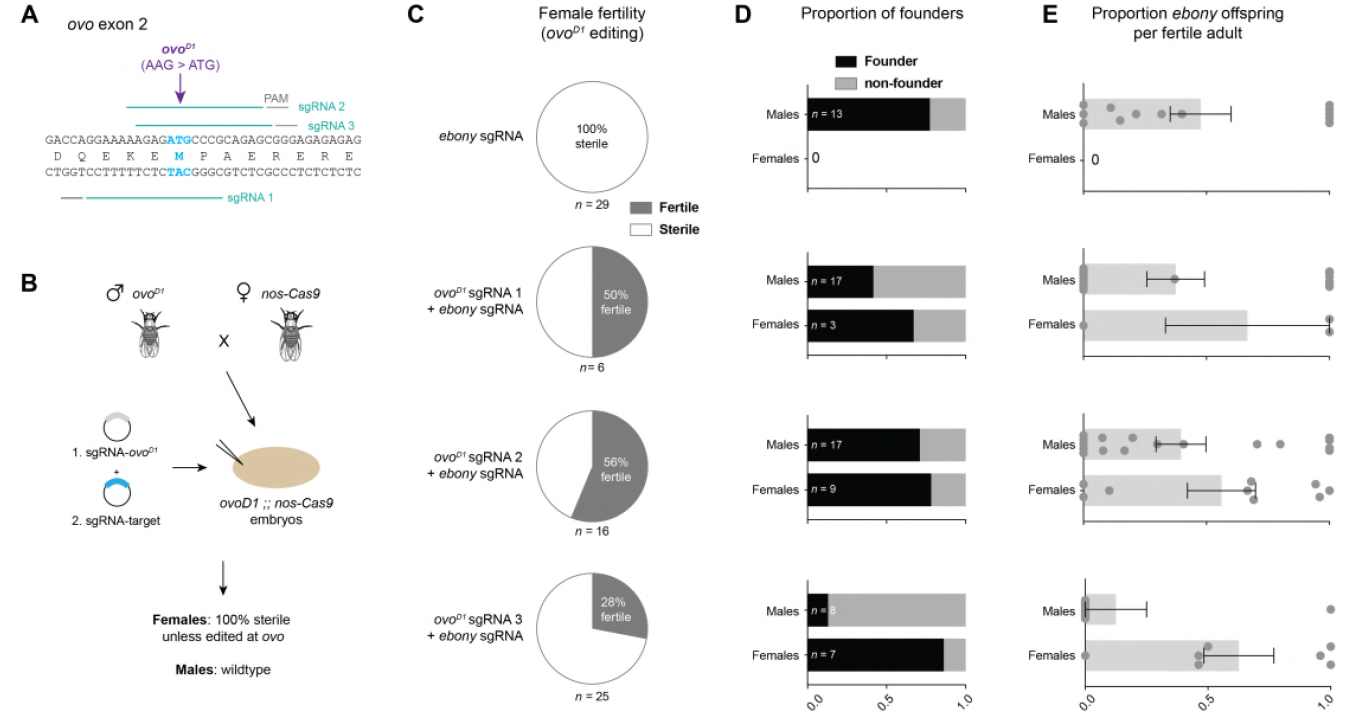
Proof of principle for *ovo^D^* co-selection. (A) Design of three sgRNAs targeting the *ovo^D1^* mutation (AAG > ATG) in the second exon of the *ovo* gene. (B) Schematic of *ovo^D^* co-selection. *ovo^D1^* males are crossed to *nos-Cas9* virgins, and their embryos (genotype = *ovo^D1^;; nos-Cas9*) are injected with an sgRNA targeting *ovo^D1^* mixed with a second sgRNA for a target gene-of-interest. GO females will be sterile unless edited at the *ovo^D1^* locus, while the males serve as an internal control. (C) Editing of the *ovoD1* locus restores fertility in a proportion of injected females. Fertility data for males is given in Table S3. Sample size is the total number of adult G0s screened for fertility (D) The proportion of fertile G0s giving rise to mutant offspring (“founders”) is higher amongst fertile females than amongst males. Sample sizes refer to the number of fertile G0s recovered from an injection. (E) The proportion of *ebony* offspring produced by each fertile G0. Each dot represents the proportion of *ebony* offspring generated by a single fertile G0 fly. Error bars show standard error of the mean.

We reasoned that CRISPR editing of the *ovo^D1^* mutation in the female germline should restore fertility specifically in successfully edited germs cells, and thus any eggs produced by such females will necessarily have undergone CRISPR editing at the *ovo^D1^* locus (**Figure 1B**). Thus, given the observed tendency for CRISPR events to co-occur in individual cells, this strategy should allow us to enrich for editing at a secondary site in all offspring (Kim *et al*. 2014; Arribere *et al*. 2014; Liao *et al*. 2015; Shy *et al*. 2016; Ge *et al*. 2016; Agudelo *et al*. 2017).

To test whether *ovo^D1^* editing indeed restores fertility, we designed three sgRNAs targeting the *ovo^D1^* locus (**Figure 1A, Table S1**). We crossed *ovo^D1^* males to *nos-Cas9* females to generate *ovo^D1^;; nos-Cas9* embryos (**Table S2** gives all *Drosophila* genotypes), and in three separate experiments, injected each of the three ovo^D1^-sgRNAs, along with an sgRNA targeting a secondary gene, *ebony*. Once mature, these injected G0 flies were individually mated, and screened for fertility. We confirmed complete sterility of *ovo^D1^;; nos-Cas9* females in uninjected controls (*n* = 3 independent crosses, 10 females per cross), consistent with previous observations (Busson *et al*. 1983). Similarly, female *ovo^D1^;; nos-Cas9* embryos injected with sgRNA-ebony alone were 100% sterile, as expected (**Figure 1C**). However, injection of any of the three sgRNA-*ovo^D1^* plasmids led to a restoration of fertility in a portion of injected females (28% - 56%, **Figure 1C, Table S3**), indicating that editing of *ovo^D1^* had occurred in a subset of germ cells. Thus, CRISPR editing of *ovo^D1^* can indeed restore germ cell function in females.

We note that a number of different editing events could conceivably restore wildtype *ovo* function, including inframe deletions that remove the novel methionine, or frameshift mutations that introduce a premature stop in the mutant allele, as females heterozygous for *ovo* loss-of-function mutations are fertile. In addition, because the wildtype and mutant forms of *ovo* differ by only one SNP, it is possible that sgRNAs targeting *ovo^D1^* form may also cleave the wildtype copy in some cases. However, any editing events that do not leave at least one wildtype copy of *ovo* intact will never be observed in offspring.

### Co-selection with *ovo^D1^* enriches for independent knock-out events at an unlinked site

To test whether *ovo^D1^* enriches for editing at a secondary locus, we scored the offspring of all fertile G0 females (i.e. those that had been edited at the *ovo^D1^* locus) for editing at a second site, *ebony*, for which we had co-injected an additional sgRNA. We screened for *ebony* knock-out alleles via complementation tests with a known allele of *ebony* (see Methods). As an internal control for each injection, we used the proportion of *ebony* alleles generated by male G0 flies, as their fertility is unaffected by *ovo^D^* (Busson *et al*. 1983). In separate control experiments, we confirmed that the frequency of CRISPR mutations for *ebony* do not differ between male and female G0s (**Figure S1A,B,C**).

For all three sgRNA-*ovo^D1^* constructs, we observed an enrichment of *ebony* editing in females compared to males (**Figure 1D,E**). The enrichment achieved by *ovo^D^* co-selection manifested in two related ways. First, the proportion of fertile G0 females giving rise to *ebony* offspring (which we refer to as “founders”) was always higher than the proportion of founders observed amongst male G0s (**Figure 1E**). Second, the average number of *ebony* offspring produced by fertile G0 females was consistently higher than produced by males (**Figure 1D**). We note that the proportion of male founders (i.e. internal controls for each injection) with successful *ebony* editing in their germ line varied widely between injections, from 12.5% to 77% (Figure 1E), indicating stochastic variation between individual injections. In contrast, the relatively higher proportion of founders obtained via ovo^D^-selection remained consistently high between all experiments, ranging from 67% to 86% (**Figure 1E**). Thus, when using *ovo^D^* co-selection, the large majority of all injected G0 females contained germ cells with mutant alleles of a second site, thus reducing the amount screening required to recover mutants. In all subsequent experiments, we used sgRNA-*ovo^D1^*-2, as it led to the highest proportion of fertile female G0s in our pilot experiment, hereafter referred to as “sgRNA-*ovo^D1^*” (**Figure 1, Table S1**).

In many cases, researchers may wish to create an allelic series of multiple independent mutations of a given target gene. We reasoned that independent *ebony* editing events may occur in different germ cells within an individual G0 female. To test this, we sequenced multiple individual offspring from each of four fertile G0 females. In all cases, we observed multiple alleles produced by each G0 female, indicating that individual primordial germ cells within a single G0 female are independently edited at the *ebony* locus (**Figure 2**). Thus, *ovo^D^* co-selection strategy allows for multiple independent mutations to be recovered from as few as one G0 female.

**Figure 2.**
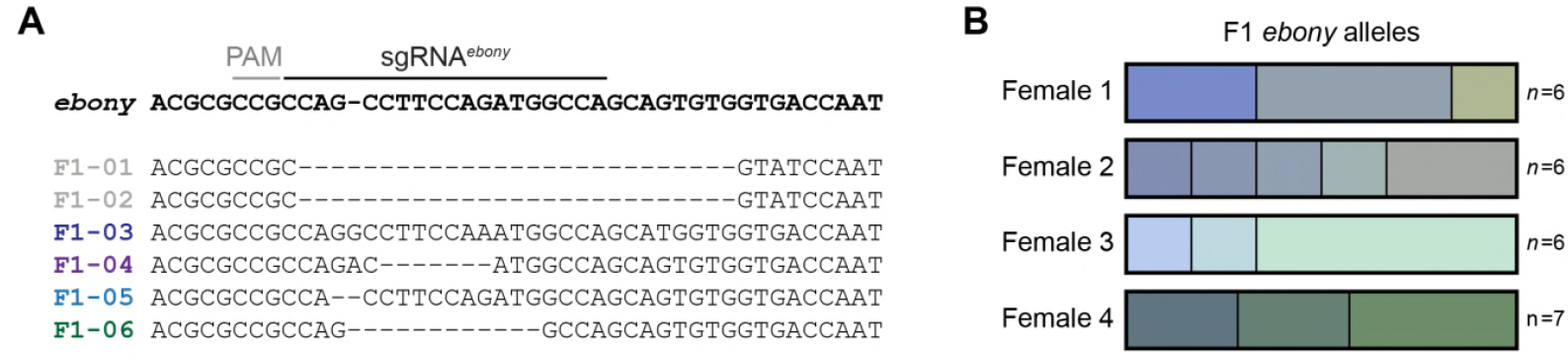
Allelic series of *ebony* generated from single fertile females using *ovo^D^* co-selection. (A) Sequence alignment of *ebony* alleles from six independent F1 generated by a single fertile female G0, indicating five separate editing events. (B) Diagram of independent *ebony* alleles identified from four individual fertile G0s. Sample size refers to number of F1 offspring sequenced per G0. Female 2 corresponds to the sequence analysis shown in (A).

Next, we tested whether *ovo^D^* co-selection reliably enriches for secondary CRISPR events by performing three additional *ovo^D^* co-selection experiments. For these experiments, we used three additional sgRNAs targeting *ebony* (Port *et al*. 2015). In all three cases, fertility was restored in between 61% - 75% of females (n = 13-16), these fertile females were enriched for founders, and their offspring were enriched for edited *ebony* alleles (**Figure 3**). Importantly, *ovoD* co-selection successfully enriched for founders regardless of the baseline effectiveness of the individual *ebony* sgRNA. For example, while sgRNA-ebony^2^ was relatively inefficient, the *ovo^D^* co-selection still enhanced the proportion of founders to from 44% in control males to 64% in females, thus reducing the amount of screening that would be necessary to obtain mutants (**Figure 3**). Thus, for each of the four sgRNAs tested, *ovoD* co-selection successfully enriches for CRISPR editing at the target site.

**Figure 3.**
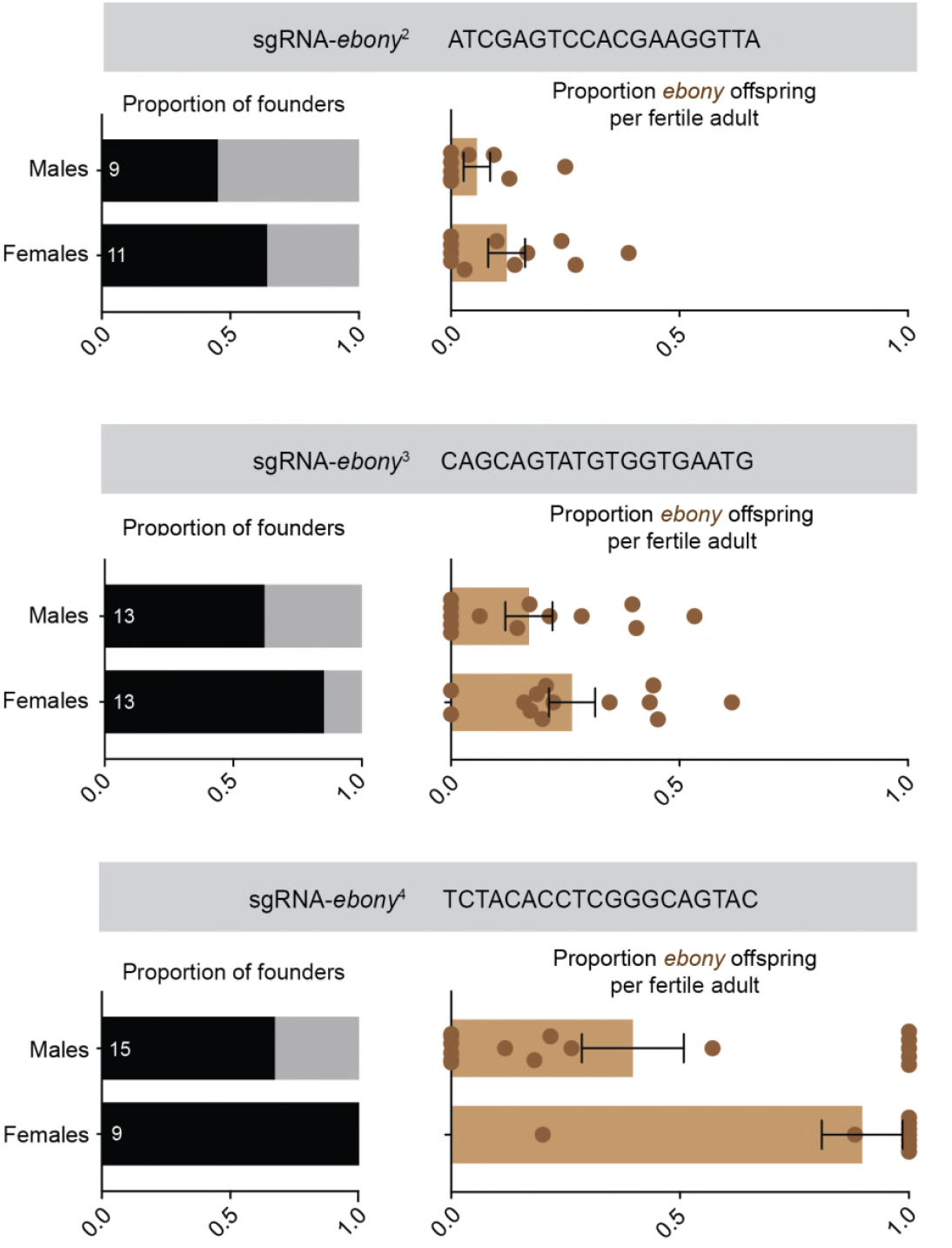
*ovo^D^* co-selection reliably enriches for CRISPR knock-out mutation events. Three independent *ovo^D^* co-selection experiments, each using a separate *ebony* sgRNA, demonstrate an enrichment of founders giving rise to edited *ebony* offspring, and an enrichment of edited offspring per founder. Sample sizes reflect the total number of adult G0s obtained for each experiment, with founders and non-founders colored as in Figure 1. Error bars show standard error of the mean.

### *ovo^D^* co-selection enriches for knock-ins

We next wished to test whether *ovo^D^* co-selection can also enrich for HDR-mediated knock-in mutagenesis. An individual cell’s propensity to repair DNA lesions via NHEJ or HDR is largely dictated by the phase of the cell cycle, with HDR largely restricted to late S/G2 phase (Heyer *et al*. 2010). Thus, in cell culture systems, it is a major challenge to enrich for CRISPR knock-in events because, at the population level, only a small minority of cells are in S/G2 at any given time, and thus NHEJ is highly favored (Agudelo *et al*. 2017). However, we noted that embryonic germ cells of *Drosophila* are arrested in G2 throughout embryogenesis (Su *et al*. 1998), suggesting that it may be possible to obtain high levels of HDR-mediated CRISPR knock-ins using our *ovo^D^* co-selection method.

To test whether *ovo^D^* co-selection enriches for knock-ins, we co-injected sgRNA-*ovo^D1^* and an sgRNA targeting an intron of *gsb-n*, together with a donor containing homology arms for *gsb-n* and a T2A-Gal4 CRIMIC insert, marked with 3XP3-GFP, a fluorescent eye marker (Lee *et al*. 2018), into *ovo^D1^;; nos:Cas9* embryos (**Figure 4A**). Fertility was restored in seven of 13 (54%) of females, of which five (71%) were founders giving rise to GFP+ offspring, compared to 38% of male G0s (**Figure 4B**). In addition, the average number of GFP+ offspring was enriched amongst female founders compared to males (**Figure 4B**.) Thus, *ovo^D^* co-selection successfully enriched for HDR-mediated CRISPR knock-in. In a separate control experiment, we injected the sgRNA and donor targeting *gsb-n* into *nos:Cas9* embryos, and confirmed that the number of founders and GFP+ offspring are equivalent in males and females (**Figure S1D**.)

**Figure 4.**
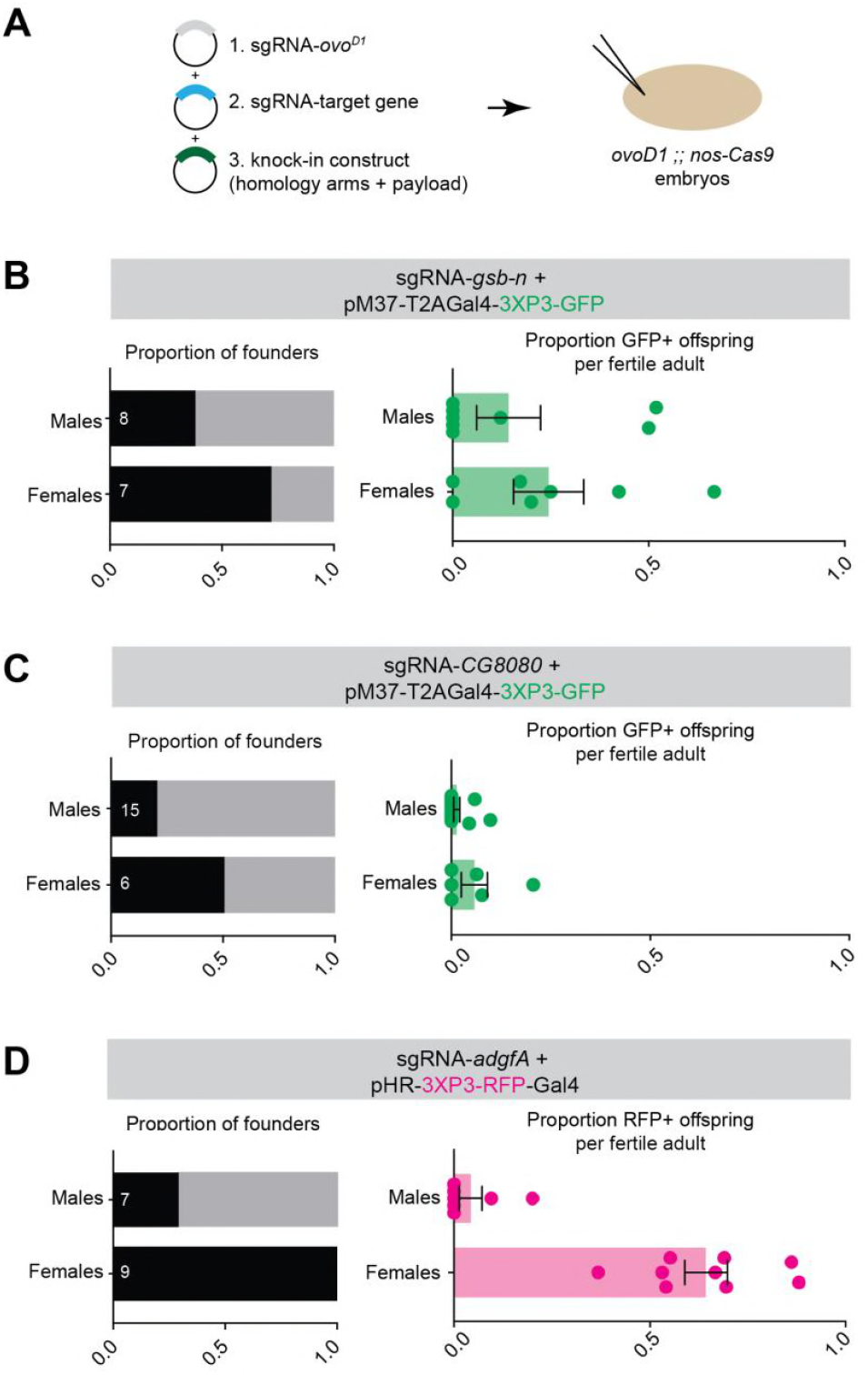
Enrichment for HDR-mediated knock-in mutagenesis using *ovo^D^* co-selection. (A) Diagram of strategy for knock-in mutagenesis using *ovo^D^* co-selection. (B-D) Three independent co-selection experiments demonstrating successful enrichment for three separate genes. Sample sizes reflect the total number of adult G0s obtained for each experiment, with founders and non-founders colored as in Figure 1. Error bars show standard error of the mean.

We repeated *ovo^D^* co-selection for two additional knock-in constructs, targeting *CG8080* and *adgf-A* with two similar donor constructs (pM37-T2A-Gal4-3XP3-GFP and pHR-3XP3-RFP, respectively). In both cases, fertility was restored in 38% - 60% (*n* = 15-16) of females, and such fertile females were enriched for founders, and their offspring were enriched for knock-in chromosomes (**Figure 4C and 4D**). We note that we observed successful enrichment in all cases despite the fact that these reagents appear to represent a range of efficiencies, with targeting of *adgf-A* being remarkably effective, and CG8080 far less so. Thus, our data suggest that *ovo^D^* co-selection reliably enriches for HDR-mediated CRISPR knock-ins as well as knock-outs.

Across all of the experiments we have conducted (*n* = nine *ovo^D^* co-selection injections), *ovo^D^* co-selection increased the proportion of successful founders by an average 35.2% (paired t-test; t=4.685, df=8, p=0.0016; **Figure 5A**). In addition, the average proportion of successful founders amongst fertile females was 77.7%, and never dropped below 50% (**Figure 5A**). In comparison, the mean proportion of founders amongst control males was 42.5%, and ranged between 12.5% - 70.5% (**Figure 5A**). *ovo^D^* co-selection also led to a 26.3% increase in the proportion of edited offspring obtained from fertile G0s compared to control males (paired t-test; t=3.623, df=8, p = 0.0068; **Figure 5B**).

**Figure 5.**
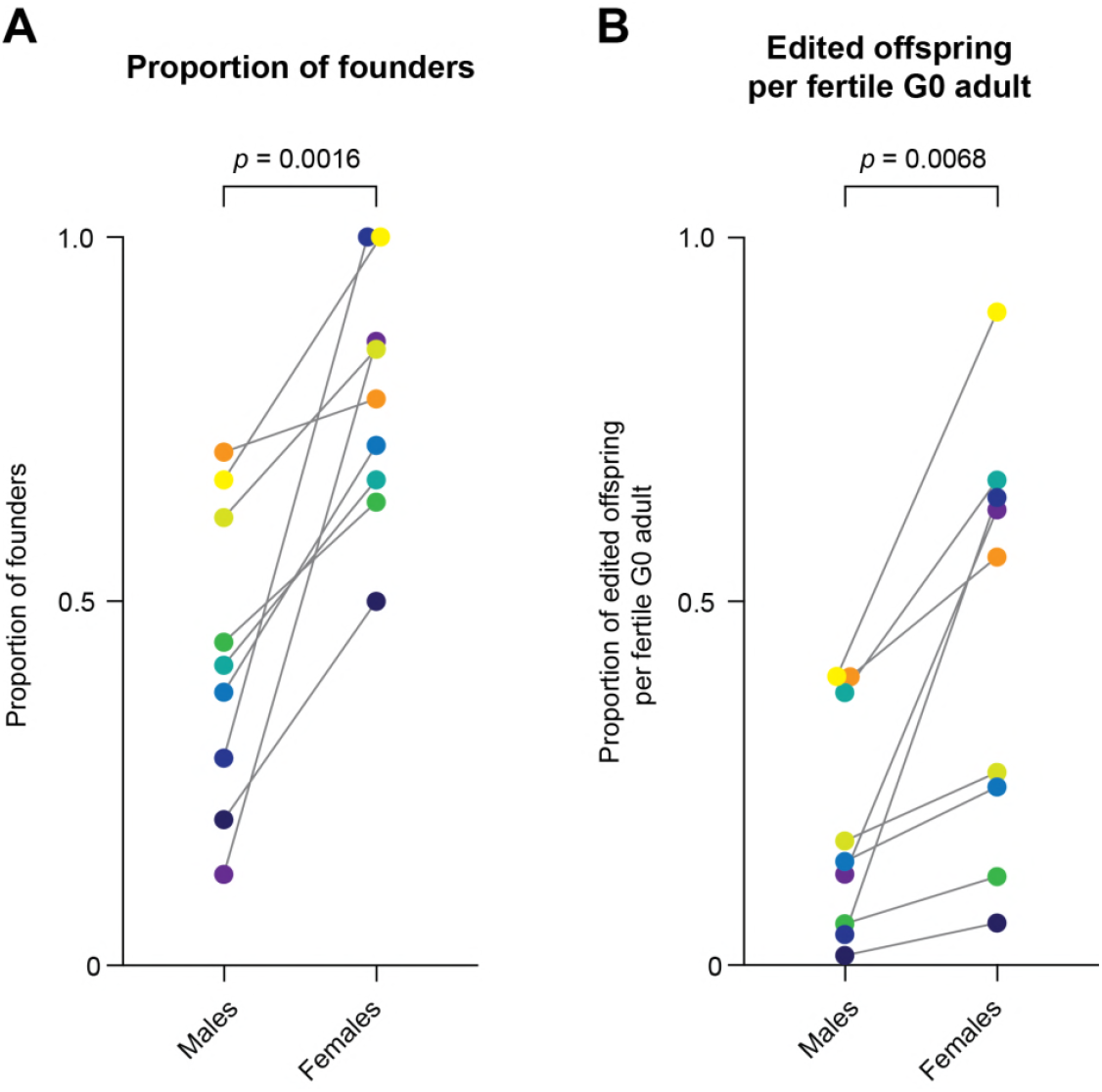
Summary of *ovo^D^* co-selection enrichment across the nine separate experiments shown in this study. (A) The proportion of founders observed amongst fertile female G0s and male G0s across all experiments shown in this study, including both knock-outs and knock-ins. Note that females represent *ovoD* co-selection, whereas males represent internal controls for each round of injection. Each dot represents one co-selection experiment, color-coded for visual clarity. *ovoD* co-selection causes a consistent and statistically significant increase in the proportion of founders. (B) The average proportion of edited offspring generated per fertile G0 is consistently and significantly enriched by *ovoD* co-selection. Colors are consistent between panels, reflecting individual injection experiments. p-values are reported for paired t-tests.

### Conclusions

A recent study of *ebony* co-selection in *Drosophila* concluded that the highest levels of CRISPR enrichment are obtained in so-called “jackpot” lines, which are those flies giving rise to very high proportions of *ebony*-offspring (Kane *et al*. 2017). Our results suggest that, using *ovo^D^* co-selection, nearly every fertile female is a jackpot line. Using this technique, the only eggs produced are those that have been edited at a minimum of one locus, which leads to a substantial enrichment of a secondary CRISPR event, both NHEJ-mediated knock-outs and HDR-mediated knock-ins. Thus, *ovo^D^* co-selection should greatly speed the recovery of CRISPR mutants, as the majority of fertile females obtained should give at least some proportion of edited offspring. As a case in point, in one of our experiments, we only obtained three fertile females from an injection, yet two of these fertile females were successful founders giving rise to high proportions of edited offspring (**Figure 1D,E**).

We propose that the mechanism of co-CRISPR enrichment is simply the successful delivery of sgRNAs and Cas9 to embryonic germ cells in a physiologically acceptable stoichiometry, and thus represents a sum total of both technical and biological variables in a given experiment. In other words, *ovo^D^* co-selection does not increase the number of CRISPR events that occur, but simply makes invisible all of the unedited germ cells, and thereby reduces the number of offspring to be screened.

The fly stocks required to perform *ovo^D^* co-selection are described in **Table S2**, and are available from the Perrimon Lab and/or the Bloomington Stock Center. The sgRNA-*ovo^D1^* plasmid is available Addgene (Plasmid 111142). In addition, we note that the there are multiple *ovo^D1^* stocks covering the second and third chromosomes, as well as germ-line specific Cas9 stocks on additional chromosomes for researchers wishing to perform CRISPR/Cas9 editing on a clean X or III chromosome.

## Acknowledgements

We thank Christians Villalta for performing injections, Charles Xu, Justin Bosch, and the Transgenic RNAi Research Project at Harvard Medical School for providing knock-in constructs, and Rich Binari for assistance with fly work. This work was supported by National Institutes of Health grant R01GM084947, and B.E.C. received National Institutes of Health funding under the Ruth L. Kirschstein National Research Service Award F32GM113395 from the NIH General Medical Sciences Division. N. Perrimon is an investigator of the Howard Hughes Medical Institute.

---

**Figure S1.**
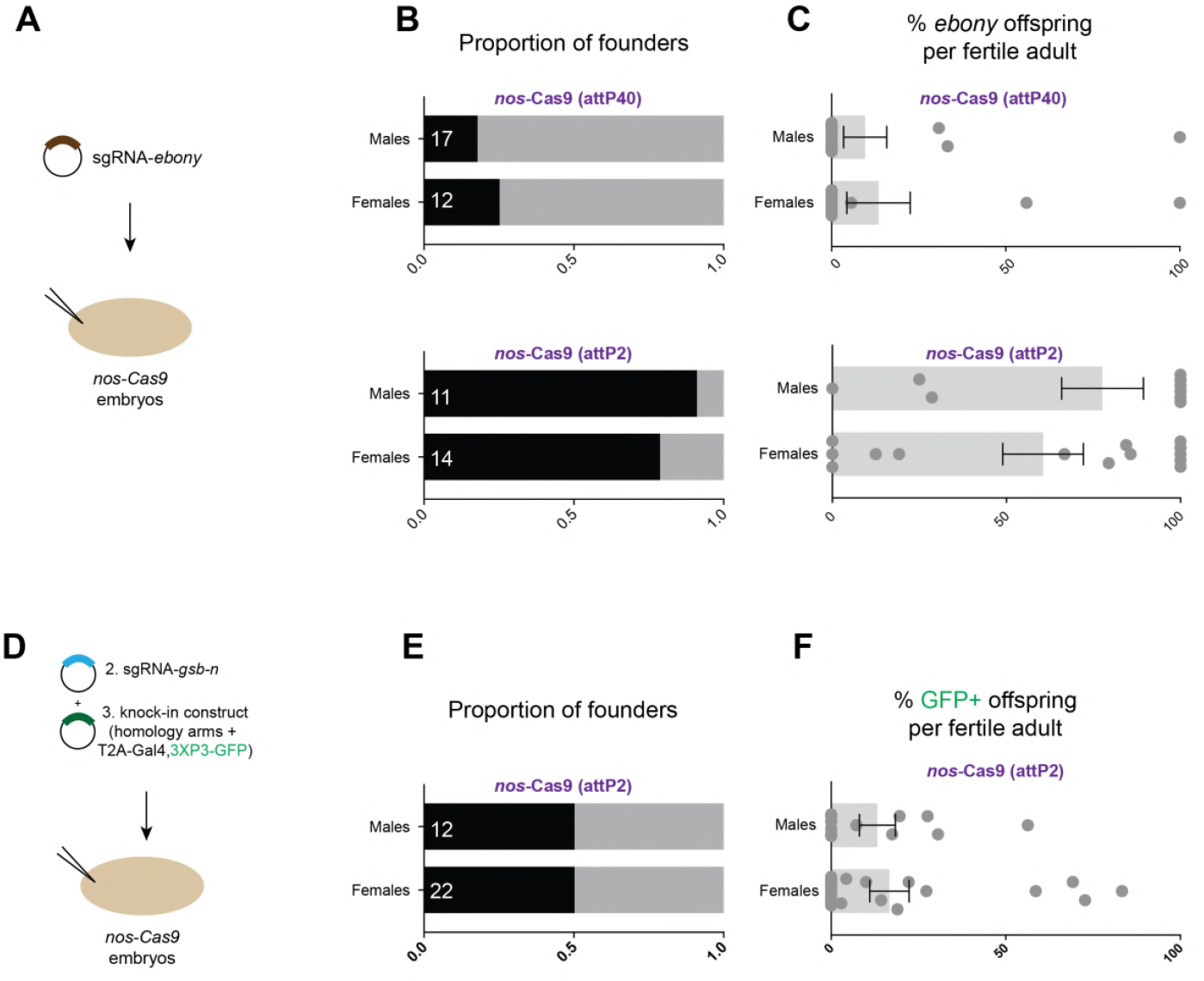
CRISPR mutagenesis is not inherently sex-biased in *Drosophila*. (A) Schematic of a standard CRISPR mutagenesis experiment, absent *ovo^D^* co-selection, in which an sgRNA targeting *ebony* is injected into a *nos-Cas9* embryo. For two separate *nos-Cas9* lines (in the attP40 and attP2 landing sites, respectively), the proportion of founders (B) and *ebony* offspring per founder do not female-biased. Note that the attP2 nos-Cas9 line is used in all other experiments in this study. (C-F) Control knock-in experiment, using the same sgRNA and donor shown in Figure 4B, but absent *ovo^D^* co-selection.

**Supplemental Table 1.**
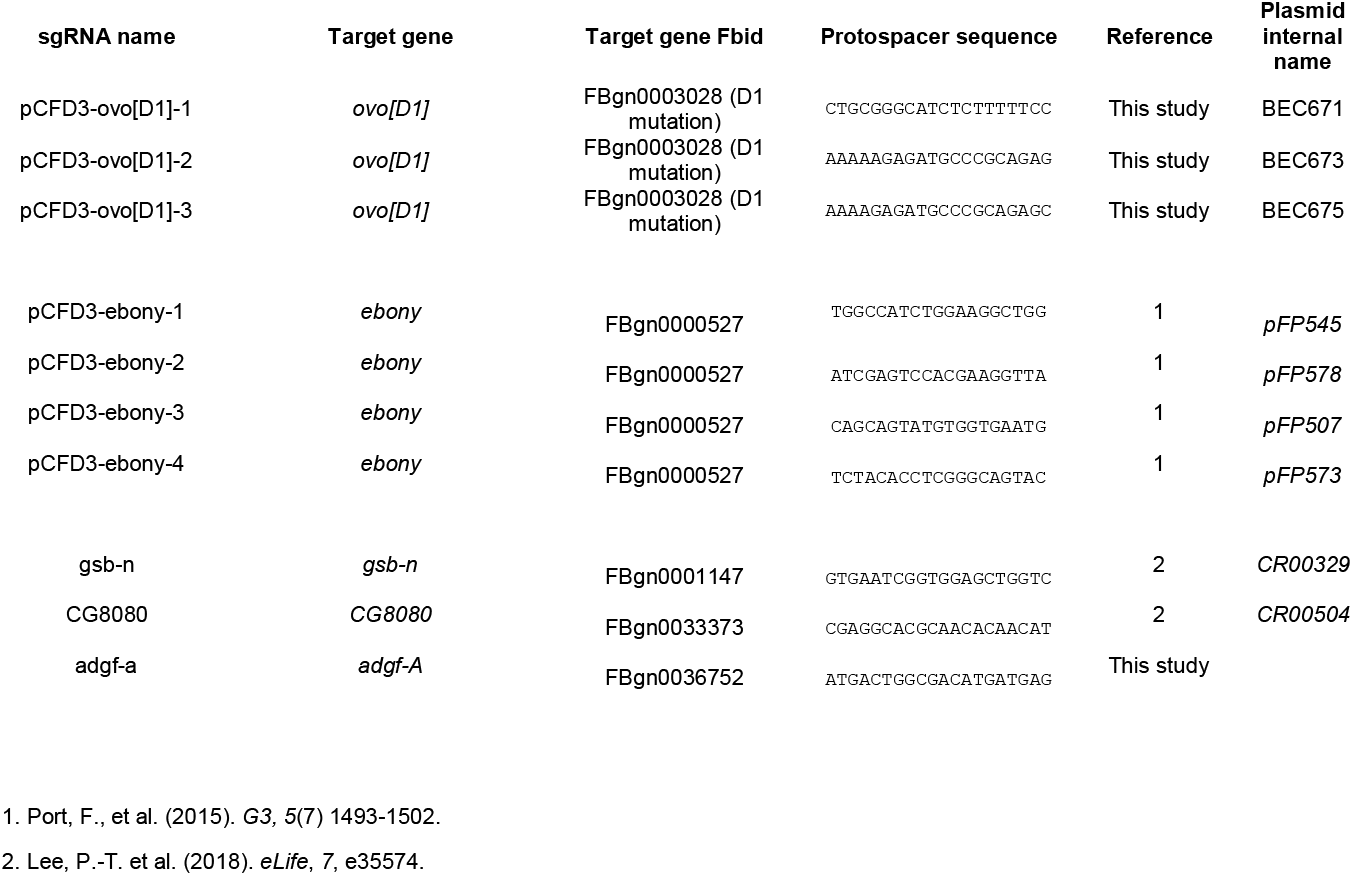
sgRNAs used in this study

**Supplemental Table 2.** *Drosophila* lines used in this study and additional *ovo^D^* stocks.

| Fly stock | Genotype | Reference | Source | Bloomington |
| --- | --- | --- | --- | --- |
| *ovoD1* (K1237) | *ovo[D1] v[24]/C(1)DX, y[1] w[1] f[1]/Y* | 1 | Perrimon Lab | 1309 |
| *yv* ;; *nos-Cas9[attP2]* | *y[1] sc[\*] v[1]; P{Nos-Cas9}Attp2* | 2 | Perrimon Lab | In progress |
| *w* ; *nos-Cas9[attP40]* | *y[1] sc[\*] v[1]; P{Nos-Cas9}Attp40* | 2 | Perrimon Lab | In progress |
| **Autosomal *ovoD* stocks not tested in this study** |  |  |  |  |
| Fly stock | Genotype | Reference | Source | Bloomington |
| *ovoD1* on 2L | *w/Y; P[ovoD1]/CyO males crossed with S Sp Ms(2)M bwD/CyO females.* | 3 | Perrimon Lab | 2121 |
| *ovoD1* on 2R | *w/Y; P[ovoD1]/CyO males crossed with S Sp Ms(2)M bwD/CyO females.* | 3 | Perrimon Lab | 2125 |
| *ovoD1* on 3L | *w/Y; P[ovoD1]/TM3, Sb males crossed with ru h st B2tD ss es/TM3, Sb females* | 3 | Perrimon Lab | 2139 |
| *ovoD1* on 3R | *w/Y; P[ovoD1]/TM3, Sb males crossed with ru h st B2tD ss es/TM3, Sb females* | 3 | Perrimon Lab | 2149 |

1. Busson, D., et al. (1983). *Genetics*, 105(2), 309–325. 2. Ren, X., et al. (2013). *PNAS*, 110(47), 19012–19017 3. Chou, T. B., & Perrimon, N. (1996). *Genetics*, 144(4), 1673–1679.

**Supplemental Table 3.** Summary statistics for each ovoD co-selection experiment.

| Testing three ovoD sgRNAs |  |  |  |  | Fertility |  | Founders |  | Mutant offspring |  |
| --- | --- | --- | --- | --- | --- | --- | --- | --- | --- | --- |
| Experiment | Genotype injected | ovoD sgRNA | target gene sgRNA | target:ovoD sgRNA ratio | % Fertile Females (n) | % Fertile Males (n) | % Female Founders | % Male Founders | Mean % Edited offspring (from female G0s) | Mean % Edited offspring (from males G0s) |
| Standard CRISPR control (attP40) | yv ;; nos:Cas9(attP40) | n/a | *ebony-1* | n/a | 92% (n = 13) | 94% (n = 18) | 25% (n = 12) | 18% (n = 17) | 9.65% | 13.47% |
| Standard CRISPR control (attP2) | yv ;; nos:Cas9(attP2) | n/a | *ebony-1* | n/a | 74% (n = 19) | 69% (n = 16) | 79% (n = 14) | 91% (n = 11) | 60.56% | 77.6% |
| no ovoD-sRNA control | ovoD1 ;; nos:Cas9 | n/a | *ebony-1* | n/a | 0% (n = 29) | 68% (n = 19) | 0 | 77% (n = 13) | - | 47.63% |
| ovoD-sgRNA 1 + ebony | ovoD1 ;; nos:Cas9 | ovoD1-1 | *ebony-1* | 1:1 | 50% (n = 6) | 100% (n = 17) | 67% (n = 3) | 41% (n = 17) | 66.67% | 37.48% |
| ovoD-sgRNA-2 + ebony | ovoD1 ;; nos:Cas9 | ovoD1-2 | *ebony-1* | 1:1 | 56% (n = 16) | 100% (n = 17) | 78% (n = 9) | 71% (n = 17) | 56.11% | 39.63% |
| ovoD-sgRNA-3 + ebony | ovoD1 ;; nos:Cas9 | ovoD1-3 | *ebony-1* | 1:1 | 28% (n = 25) | 89% (n = 9) | 86% (n = 7) | 13% (n = 8) | 62.60% | 12.50% |

| Three additional *ebony* knock-outs |  |  |  |  | Fertility |  | Founders |  | Mutant offspring |  |
| --- | --- | --- | --- | --- | --- | --- | --- | --- | --- | --- |
| Experiment | Genotype injected | ovoD sgRNA | target gene sgRNA | target : ovoD sgRNA ratio | % Fertile Females (n) | % Fertile Males (n) | % Female Founders | % Male Founders | Mean % Edited offspring (from female G0s) | Mean % Edited offspring (from males G0s) |
| *ebony-2* | ovoD1 ;; nos:Cas9 | ovoD1-2 | *ebony-2* | 2:1 | 61% (n = 18) | 69% (n = 13) | 64% (n = 11) | 44% (n = 9) | 12.19% | 5.68% |
| *ebony-3* | ovoD1 ;; nos:Cas9 | ovoD1-2 | *ebony-3* | 2:1 | 62% (n = 21) | 100% (n = 13) | 85% (n = 13) | 62% (n = 13) | 26.52% | 17.06% |
| *ebony-4* | ovoD1 ;; nos:Cas9 | ovoD1-2 | *ebony-4* | 2:1 | 75% (n = 12) | 100% (n = 16) | 100% (n = 9) | 67% (n = 15) | 89.80% | 39.69% |

| Knock-in experiments |  |  |  |  | Fertility |  | Founders |  | Mutant offspring |  |
| --- | --- | --- | --- | --- | --- | --- | --- | --- | --- | --- |
| Injection | Genotype injected | ovoD sgRNA | target gene sgRNA | target : ovoD : donor sgRNA ratio | % Fertile Females (n) | % Fertile Males (n) | % Female Founders | % Male Founders | Mean % Edited offspring (from female G0s) | Mean % Edited offspring (from males G0s) |
| Control - gsb-n CRIMIC | yv ;; nos:Cas9 | n/a | *gsb-n* | 2:-:3 | 79% (n = 28) | 75% (n = 16) | 50% (n = 22) | 50% (n = 12) | 16.69% | 13.23% |
| gsb-n CRIMIC | ovoD1 ;; nos:Cas9 | ovoD1-2 | *gsb-n* | 2:1:3 | 54% (n = 13) | 100% (n = 9) | 71% (n = 7) | 38% (n = 8) | 24.47% | 14.26% |
| adgf-a CRIMIC | ovoD1 ;; nos:Cas9 | ovoD1-2 | *adgf-A* | 2:1:3 | 60% (n = 15) | 86% (n = 8) | 100% (n = 9) | 29% (n = 7) | 64.27% | 4.21% |
| CG8080 CRIMIC | ovoD1 ;; nos:Cas9 | ovoD1-2 | *CG8080* | 2:1:3 | 38% (n=16) | 100% (n = 15) | 50% (n = 6) | 20% (n = 15) | 5.78% | 1.35% |

